# Accelerating computational Bayesian inference for stochastic biochemical reaction network models using multilevel Monte Carlo sampling

**DOI:** 10.1101/064170

**Authors:** David J. Warne, Ruth E. Baker, Matthew J. Simpson

## Abstract

Investigating the behavior of stochastic models of biochemical reactionnetworks generally relies upon numerical stochastic simulation methods to generate many realizations of the model. For many practical applications, such numerical simulation can be computationally expensive. The statistical inference of reaction rate parameters based on observed data is, however, a significantly greater computational challenge; often relying upon likelihood-free methods such as approximate Bayesian computation, that require the generation of millions of individual stochastic realizations. In this study, we investigate a new approach to computational inference, based on multilevel Monte Carlo sampling: we approximate the posterior cumulative distribution function through a combination of model samples taken over a range of acceptance thresholds. We demonstrate this approach using a variety of discrete-state, continuous-time Markov models of biochemical reactionnetworks. Results show that a computational gain over standard rejection schemes of up to an order of magnitude is achievable without significant loss in estimator accuracy.

**Author Summary:** We develop a new method to infer the reaction rate parameters for stochastic models of biochemical reaction networks. Standard computational approaches, based on numerical simulations, are often used to estimate parameters. These computational approaches, however, are extremely expensive, potentially requiring millions of simulations. To alleviate this issue, we apply a different method of sampling allowing us to find an optimal trade-off between performance and accuracy. Our approach is approximately one order of magnitude faster than standard methods, without significant loss in accuracy.

## Introduction

Stochastic models of biochemical reaction networks often provide a more accurate description of system dynamics than deterministic models [1]. In many cases, this is due to the inherent stochastic nature of many biochemical processes in which the system dynamics is significantly influenced by relatively low populations of certain chemical species [2]. For example, in eukaryotic cells, molecules that regulate gene expression occur in relatively low numbers; as a result, stochastic fluctuations have a direct effect on the production rates of proteins [3, 4].

A common approach to modeling biochemical systems is to consider a well-mixed collection of molecules that react according to some known chemical reactions. The well-mixed assumption simplifies the model by removing the spatial component [5, 6]. If the model is deterministic, evolution of the concentrations of each chemical species is governed by a system of ordinary differential equations (ODEs). Alternatively, a stochastic model will typically consider the evolution of copy numbers (i.e., the numbers of molecules) of each species, with each reaction occurring stochastically [6].

In the case of the stochastic model, the *probability density function* (PDF) for the state of such a system at time *t* evolves according to a very large system of ODEs known as the *chemical master equation* (CME), which is in general intractable due to the very large, or countably infinite, number of possible system states [5, 7]. As a result, stochastic simulation techniques such as the exact *Gillespie direct method* (GDM) or approximations like the *tau-leaping* method are applied to study these models [8, 9]. However, accurate stochastic simulation is a computationally expensive task; the computation time for the GDM, for example, scales with the number of possible reactions yet the performance improvements gained using approximations can introduce approximation errors. Therefore, the development of efficient and accurate stochastic simulation algorithms is an area of active research [10–16].

In order to make quantitative predictions of real biochemical systems or to perform model validation, unknown reaction rate parameters must be determined through inference. The Bayesian approach to estimate an unknown parameter vector, ***θ***, given some observational data, **𝒟**, is based on Bayes’ Theorem 
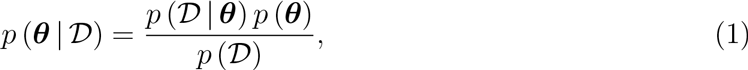
where *p* (***θ***) is the *a priori* PDF of the unknown parameter, *p* (**𝒟** ∣ ***θ***) is the likelihood of making observations **𝒟** under the assumption of a particular value for ***θ***, *p*(**𝒟**) is often referred to as the evidence and *p* (***θ*** ∣ **𝒟**) is the *a posterior* PDF of ***θ*** given the observations [17]. Informally, Equation (1) represents the process of updating current understanding based on previous experience and observational data. The classical approach to inference is to maximize the right hand side of Equation (1) to determine the mode of the posterior. However, more generally, the Bayesian approach can be used to quantify the level of uncertainty associated with such parameter estimates.

Theoretically, given perfect observational data from a biochemical reaction network, it is possible to determine a closed form expression for the likelihood term in Equation (1) and the method of maximum likelihood may be directly applied [6]. In practice, however, such a process is sampled imperfectly and is subject to measurement errors; thus requiring solution of the CME to form the likelihood term. However, as we have noted, exact closed form solutions of the CME are rarely available for practical applications.

Approximate Bayesian computation (ABC) refers to a family of computational methods for performing inference for problems with intractable likelihoods [17, 18]. As a result, ABC methods are routinely applied to practical inference problems [19–24]. The fundamental concept is to approximate the posterior PDF, *p* (***θ*** ∣**𝒟**), by *p* (***θ*** ∣ (*ρ* (**𝒟**_*s*_, **𝒟**)< ϵ), where **𝒟**_*s*_ is simulated data, *ρ* is a suitable distance function and ϵ is the acceptance threshold. If a simulation process exists for the prior distribution, *p* (***θ***), and the underlying model of interest can be simulated to estimate *p* (**𝒟** *_s_* ∣ ***θ***), then the approximated posterior can be simulated using an ABC approach.

The computational overhead for ABC inference is significantly greater than that of stochastic simulation alone. This is because the computation time is inversely proportional to the probability of *ρ* (**𝒟**_*s*_, **𝒟**) < ϵ over all possible parameter values ***θ***; that is, many stochastic simulations are required for every sample computed from the posterior [17]. A considerable computational problem arises from this, because the acceptance rate decreases exponentially as the number of unknown parameters increases [25]. This problem, referred to as the *curse of dimensionality*, is mitigated to some degree for certain classes of problems through the use of more advanced ABC techniques [26–29]. However, in general, the curse of dimensionality is an unresolved problem [17].

In this study, we propose, implement and analyze a method for accelerating ABC inference using a *multilevel Monte Carlo* (MLMC) approach [30]. MLMC is a framework for constructing computationally efficient and accurate estimators of system statistics for stochastic processes. MLMC was developed by Giles et al. [31, 32] as a stochastic variant of multigrid methods used for obtaining numerical solutions to differential equations. Since then many other applications have benefitted from MLMC including Markov process simulation [14, 33, 34], uncertainty quantification [35] and univariate probability distribution approximation [36, 37]. To the best of our knowledge, our work represents the first application of MLMC to full Bayesian inference with intractable likelihoods.

To summarize our approach, we construct an approximation to the posterior *cumulative distribution function* (CDF) using a MLMC estimator that is constructed from a sequence of ABC approximate posteriors. While we focus on stochastic biochemical reaction networks models, our inference method is applicable for any problem that could traditionally utilize ABC inference. We demonstrate that our method is guaranteed to out perform standard ABC methods under a few reasonable assumptions.

## Methods

In this section, we describe a commonly used stochastic approach to modeling biochemical reaction networks along with standard algorithms for both simulation and parameter inference. We also describe the fundamental concepts of MLMC. Finally, we present our multilevel approach to ABC inference and derive the asymptotic performance improvement of the method.

### Discrete-state, continuous-time Markov processes

Consider a biochemical reaction network involving *N* chemical species with copy numbers *X*_1_, *X*_2_,…, *X*_*N*_ that react via a network of *M* chemical reactions of the form 
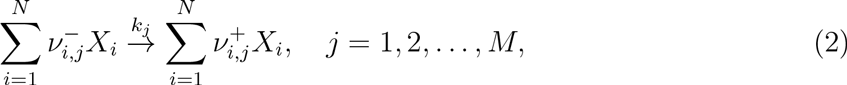
where *k*_*j*_ is the kinetic rate constant of reaction *j*, and 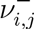 and 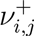 are, respectively, the reactant and product stoichiometries for the species *X_i_* in reaction *j*. Under the assumption that the molecules are well-mixed, the probability that the *j*th reaction occurs in the time interval (*t*, *t*+Δ*t*] is given by *a*_*j*_ (**X** (*t*);*k*_*j*_)Δ*t* where *a*_*j*_ is the propensity function of the reaction, and is given by 
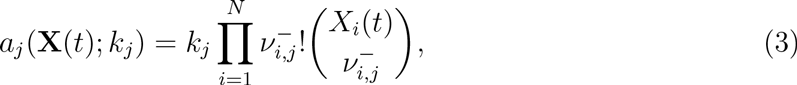
where **X**(*t*) = [*X*_1_(*t*), *X*_2_(*t*),…, *X*_*N*_(*t*)]^T^ is the column vector of copy numbers representing the system state at time *t*.

We can model the reaction network defined by Equation (2) as a discrete-state, continuous-time (DSCT) Markov process in which each reaction channel is governed by a time-varying Poisson process with rate parameter 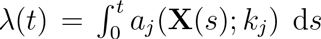. Such a DSCT Markov process can be represented according to the Kurtz representation [38] as 
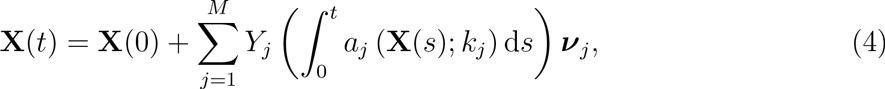
where the *Y*_*j*_(λ) are unit time Poisson processes with rate parameter λ and ***ν***_*j*_ is the state transition that results when reaction *j* takes place, that is 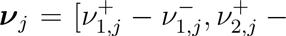 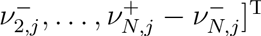.

Let *p*(*x*, *t* ∣ *y*, *s*) denote the *transitional density function* of the DSCT Markov process given in Equation (4), that is, the probability **X**(*t*) = **x** given **X**(*s*) = **y** where *t* > *s*. Given an initial condition, **X**(0) = **x**_0_, the evolution of *p*(**x**, *t* ∣ **x**_0_, 0) is governed by the CME, 
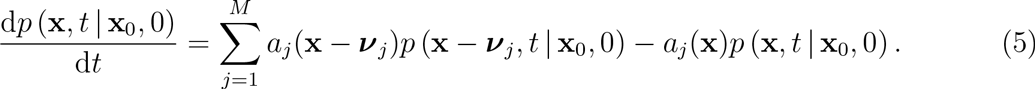

It should be noted that Equation (5) is actually a system of ODEs that is potentially countably infinite since **x** ∊ ℕ *^N^*. With the exception of reaction networks involving only zeroth and first order reactions the CME has no closed form solution, and it is generally only computationally tractable when the number of possible states is small [7, 39].

### The chemical master equation and Bayesian inference

The solution of Equation (5) is of critical importance to the Bayesian approach of parameter inference for DSCT Markov processes [6]. Given a realization of Equation (4), **X**_*d*_(*t*), observed at *N*_*t*_ discrete points in time, *t*_1_ < *t*_2_ <… < *t*_*N*_*t*__, the inference problem is to determine the posterior PDF, 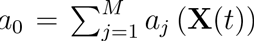, for the kinetic rate parameters, ***θ*** =[*k*_1_, *k*_2_,…, *k*_*M*_]. If we denote *p*(***θ***) as the prior PDF, that represents some prior knowledge about ***θ***, then the posterior PDF is given through application of Bayes’ Theorem (Equation (1)),

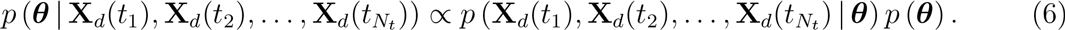

The likelihood, *p*(**X**_*d*_(*t*_1_), **X**_*d*_(*t*_2_), …, **X**_*d*_ (*t*_*N*_*t*__) ∣***θ***), can be expressed in terms of the transitional density function *p*(**X**,*t*∣***y***, *s*; ***θ***); that is, the solution to the CME parameterized by the kinetic rate parameter vector ***θ***. The likelihood term becomes, 
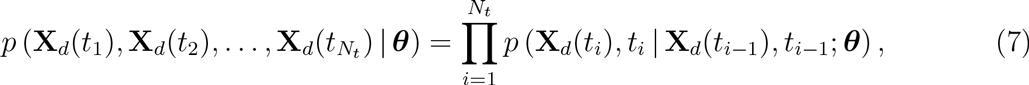
where *t*_0_ = 0 and **X**_*d*_(0) = **x**_0_.

We now introduce two models with convenient closed form solutions to the CME for theoretical purposes. Our aim in studying these two relatively simple models with closed form solutions is to illustrate the mathematical rigor of our analysis. Once we have established this, we will then apply our method to more practical examples where the CME is intractable.

### Example 1: Degradation

The simplest chemical reaction model one could conceive is the stochastic form of exponential decay; that is, the degradation model. This model has a single reaction, 
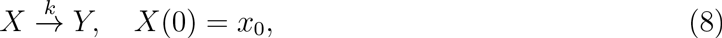
where *Y* represents any chemical species that is not of interest, *k* is the kinetic rate constant for the reaction, and **X**_0_ is the initial condition. Figure 1(a) shows some typical realizations of this model generated using the GDM; note that **X** can never increase in time.

**Figure 1.**
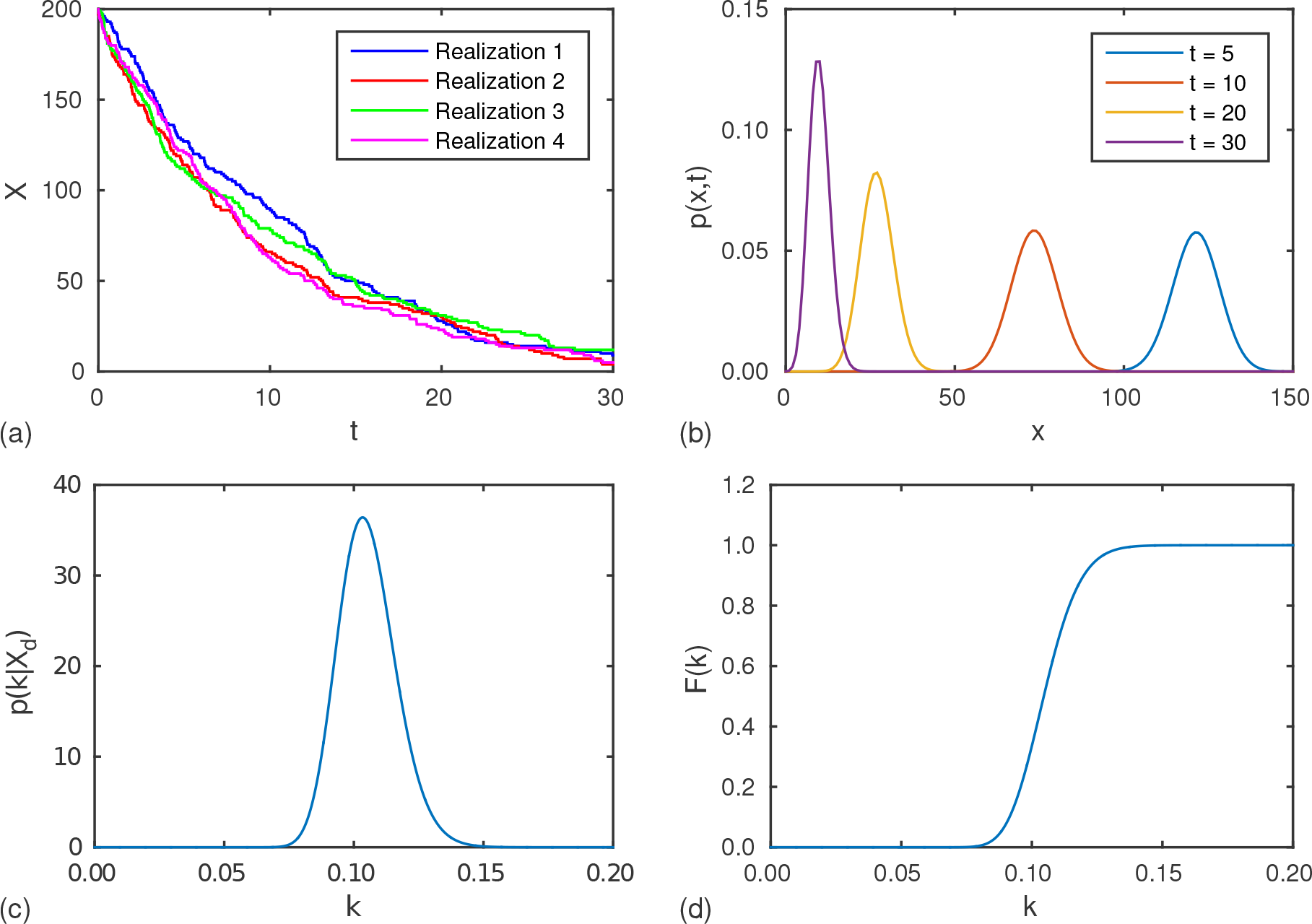
Degradation model. (a) Example realizations with *k* = 0.1 (sec^-1^) and *x*_0_ = 200. (b) Evolution of the CME solution, *p*(*x*, *t* ∣ *x*_0_, 0; *k*). The posterior (c) PDF and (d) CDF, for *X*(0) = 200 and *X*(30) = 9.

Let *p* (*x*, *t* ∣ *x*_0_, 0; *k*) be the transitional density function of the DSCT Markov process for Equation (8); that is, the solution to the CME parameterized by the kinetic rate parameter, *k*. The closed form solution to the CME can be obtained by noting that *p*(*x*, *t* ∣ *x*_0_, 0; *k*) for any *x* > *x*_0_, hence the CME can be truncated into a finite system of ODEs that may be solved through induction. The solution is given by [40] 
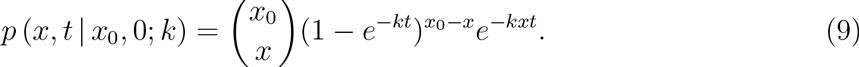

The evolution of *p* (*x*, *t* ∣ *x*_0_, 0; *k*)is shown in Figure 1(b).

For the purposes of inference of *k*, if it is given that *X*(*t*) = *x*, we can view Equation (9) as the likelihood term in Equation (1) when the number of observation times is *N*_*t*_ = 1. This enables the direct evaluation of the posterior PDF and *cumulative distribution function* (CDF) for the degradation model. Examples are given in Figure 1(c)-(d).

### Example 2: Degradation/Production

A natural extension to the degradation model (Equation (8)) is to incorporate a production reaction, 
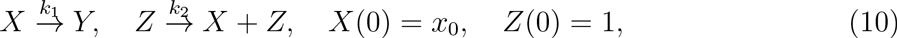
where *k*_1_ and *k*_2_ are, respectively, the degradation and production kinetic rate constants and *x*_0_ is the initial condition. Again, *Y* denotes any chemical species that is not of interest, and *Z* represents a chemical species or process that produces *X*. We note that the copy number of *Z* is constant, thus it is not required to be included in the CME. The example realizations that are shown in Figure 2(a) demonstrate the fact that *X* is not strictly decreasing in time, thus the CME cannot be truncated into a finite system of ODEs without approximation.

In this case, we denote the transitional density function as *p* (*x*, *t* ∣ *x*_0_, 0; *k*_1_, *k*_2_). Despite the countably infinite nature of the CME in this case, it can also be solved analytically [41] to give 
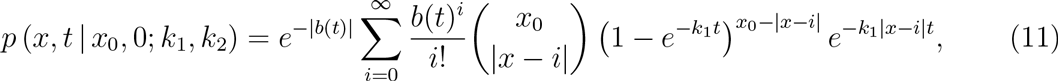
where 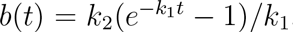 The evolution of *p* (*x*, *t* ∣ *x*_0_, 0; *k*_1_, *k*_2_) approaches a steady state by approximately *t* = 30 (sec) as shown in Figure 2(b). Just as with the degradation model, the exact posterior PDF can be derived using Equations (7) and (11) for the purposes of inference of both *k*_1_ and *k*_2_ given *X* at discrete points in time. Figure 2(c)-(d) shows examples of the posterior PDF and CDF.

**Figure 2.**
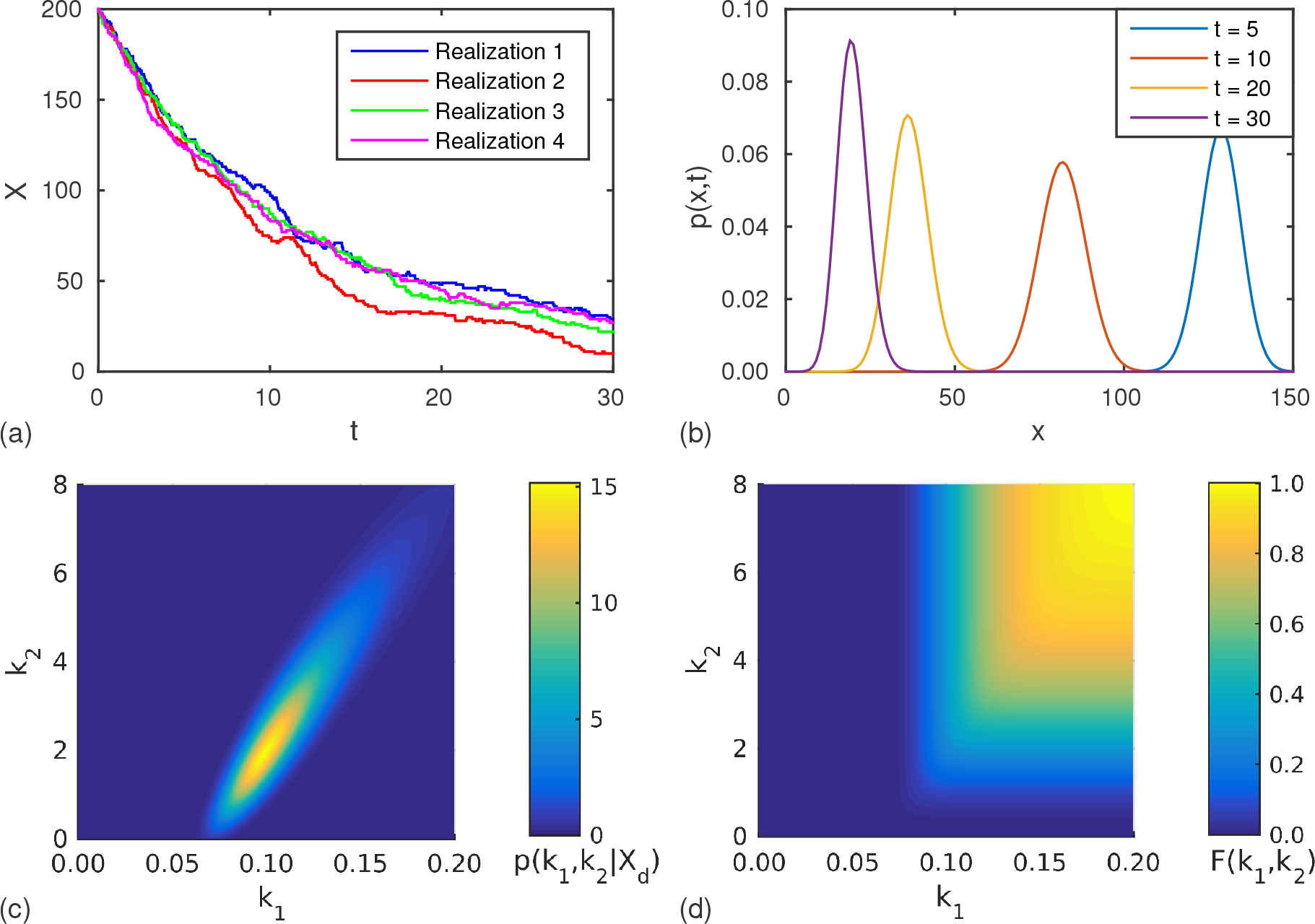
Degradation/production model. (a)Example realizations with *k*_1_ =0.1 (sec^-1^), *k*_2_ = 1.0 (sec^-1^) and *x*_0_ = 200. (b) Evolution of the CME solution, *p* (*x*, *t* ∣ *x*_0_, 0; *k*_1_, *k*_2_). The posterior (c) PDF and (d) CDF, for *X*(0) = 200, *X*(15) = 60 and *X*(30) = 29.

### Stochastic simulation: Gillespie direct method

In general, only models dealing with zeroth and first order reactions have closed form solutions [41]. If we wish to study stochastic models of chemical reaction networks that have higher order reactions, then stochastic simulation must be utilized. The most well known *exact* stochastic simulation algorithm is the GDM [8].

The GDM arises naturally from Equation (4) by recalling that the time to the next event of a Poisson process with rate parameter λ is exponentially distributed with parameter λ. Therefore at time *t*, if 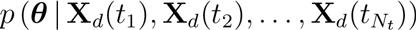, then Δ*t* ~ Exp(*a*_0_) where the next reaction occurs at *t* + Δ*t*. The next reaction, *R*, to take place can be determined by sampling the probability mass function *p* (*R* = *j*) = *a*_*j*_/*a*_0_. The state vector can then be updated by adding ***v***_*j*_. The result of repeating this process up to a given end time, *T*, is the GDM, as shown in Algorithm 1.

**Algorithm 1.**
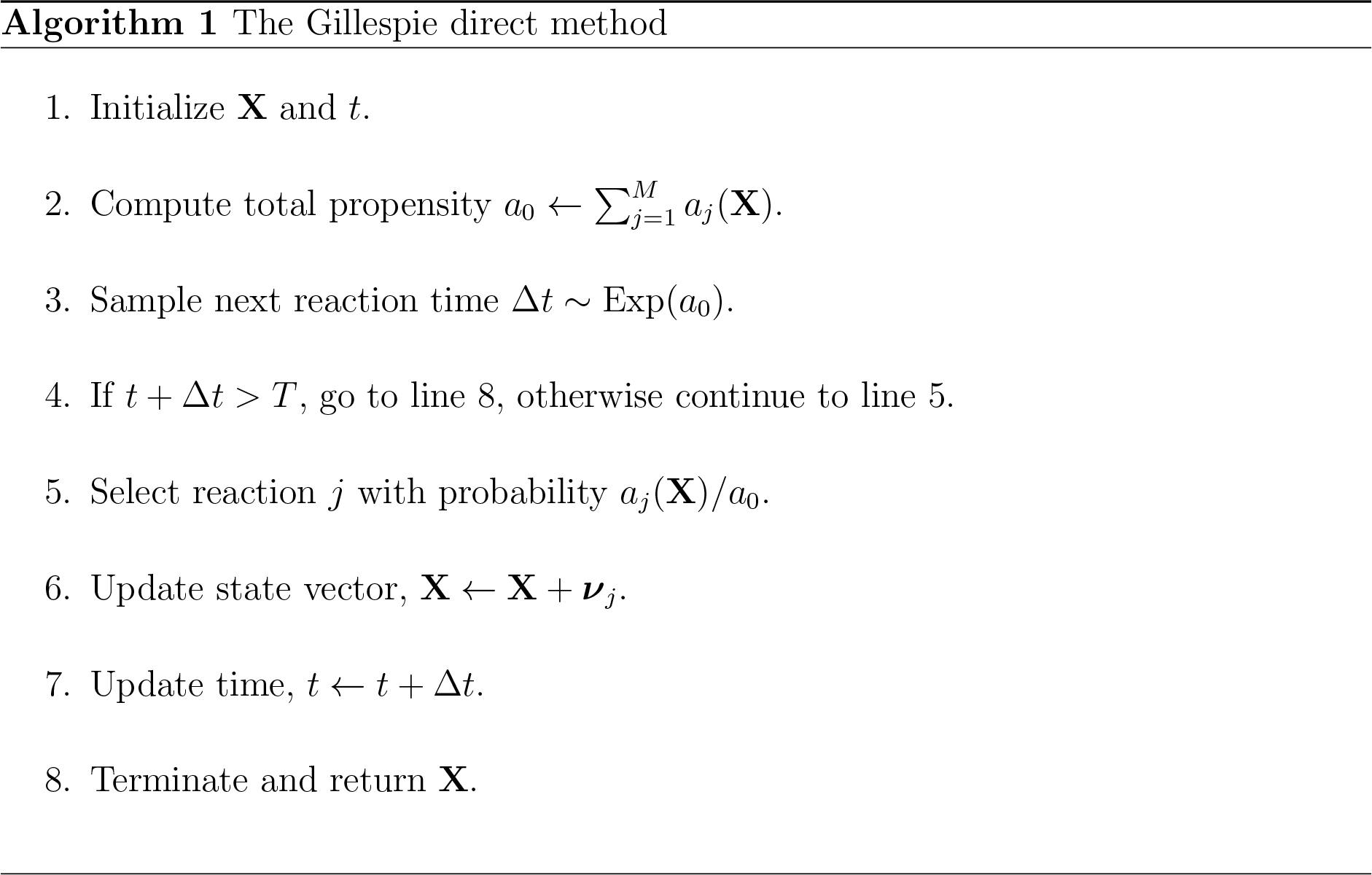

In this study, we restrict our experimentation and discussion to the GDM for stochastic simulation to ensure the only source of approximation error is due to our inference method. As a result, we do not consider approximations such as the tau-leaping method [9].

### Parameter inference: ABC rejection

ABC methods provide a means of sampling an approximation to the posterior when the stochastic process of interest has an intractable likelihood but can be simulated [6, 17]. In the context of biochemical reaction networks, this means that repeated sampling of the model using the GDM replaces the explicit solution of the CME.

Given a data set, **𝒟**, and a prior distribution for the parameter of interest, ***θ***, we approximate Equation (1) with 
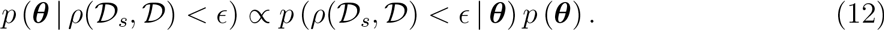

In Equation (12), **𝒟**_*s*_ is simulated data from the model of interest, *ρ* is a suitable distance metric and e is a sufficiently small acceptance threshold, such that *p* (***θ*** ∣ *ρ*(**𝒟***_s_*,**𝒟**) < ϵ) ≈ *p* (***θ*** ∣ **𝒟**). The most direct approach to sampling the posterior in Equation (12), given in Algorithm 2, is the *ABC rejection sampling* method [6, 17, 18].

**Algorithm 2.**
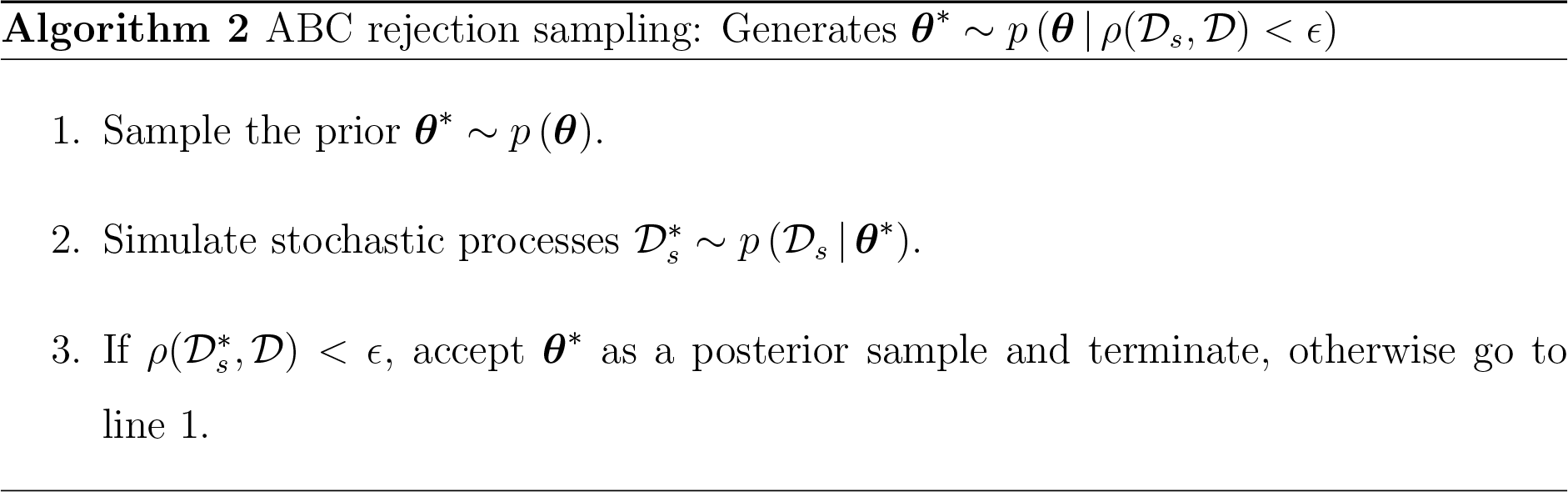

For this study, the data set, **𝒟**, for the model of interest is assumed to be the observation of a single realization, **X**_*d*_(*t*), observed at *N*_*t*_ uniformly spaced time points *t_i_* = iΔ*t* with *i* = 1, 2,…, *N*_*t*_. That is, 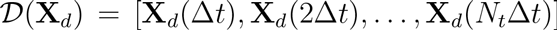. Similarly, simulated data is given by 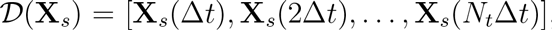, where **X**_*s*_(*t*) is a sample path generated by the GDM using the candidate parameter vector, ***θ***\*, that has been sampled from the prior distribution *p* (***θ***). A natural distance metric, *ρ*, for such data is

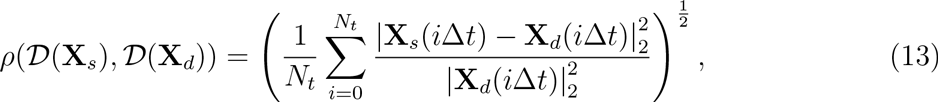
where ∣․∣_2_ is the *l*_2_-norm.

### Multilevel Monte Carlo sampling

To explain the basics of MLMC, consider the task of computing 𝔼 [*f*(*X*)] for a random variable *X* with unknown distribution and a suitably defined functional *f*. If we have another random variable *Y* that approximates *X* then we have, 
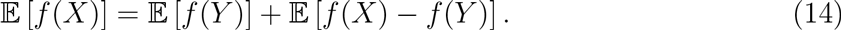

That is, an unbiased estimator for 𝔼 [*f*(*y*)] will be a biased estimator for 𝔼 [*f(X)*]. If it is possible to simulate *X*, then we can correct for this bias by using an estimator for the bias 𝔼 [*f*(*x*) – *f*(*y*)] [31]. If *Y* can be constructed in such a way that the estimator for 𝔼 [*f*(*y*)] and the bias 𝔼 [*f*(*x*) – *f*(*y*)] can be computed more efficiently than the estimator for 𝔼 [*f*(*x*)] then we have a net computational gain.

MLMC extends this idea to the case when there exists a sequence random variables 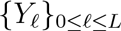 such that as *ℓ* increases the simulation time for 𝔼 [*f*(*Y*_*ℓ*_)] increases and the bias 𝔼 [*f*(*x*) – *f*(*Y*_*ℓ*_)] decreases at a suitable rate [31, 37]. The resulting recursive application of Equation (14) yields the telescoping sum 
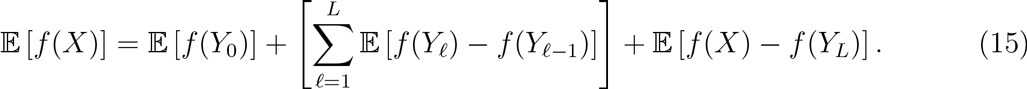

Under certain conditions, constructing an estimator based on Equation (15) is significantly faster than estimating 𝔼 [*f*(*x*)] directly [31, 33, 34].

### A multilevel approach to inference

In this section, we develop a new approach to ABC inference using MLMC and provide key theoretical results about the performance gain using our approach. The method, which we refer to as *multilevel approximate Bayesian computation* (MLABC), combines ABC rejection sampling using a sequence of acceptance thresholds to approximate the CDF of the posterior to within a prescribed level of accuracy defined in terms of the root mean squared error (RMSE). For brevity, we only present the key features of the analysis here; for detailed analysis of the method, based on the work of Giles et al. [37], see Appendices S1-S3.

### Derivation

We present, for the sake of simplicity, our MLABC method in terms of the degradation/production model (Equation (10)). However, we note that applying the ideas to different models with different numbers of unknown parameters is straightforward extension of the degradation/production method. Given observed trajectory data, **𝒟**(**X**_*d*_), the task is to approximate, at a point (*s*_1_, *s*_2_) ∊ ℝ^2^, the posterior CDF given by 
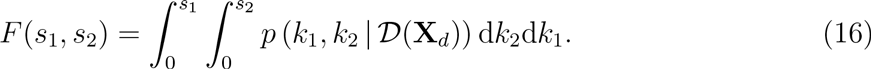

We can reformulate Equation (16) as the expectation, 
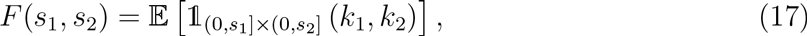
where 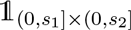 is the indicator functional, 
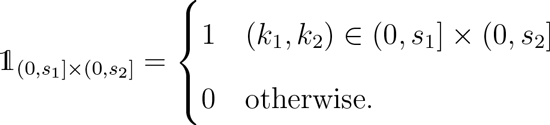

Assuming distance metric *ρ*, as defined in Equation (13), along with a suitably chosen acceptance threshold, ϵ, the standard Monte Carlo estimator is 
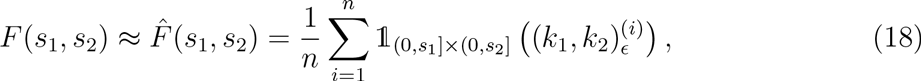
where the number of samples, *n*, is sufficiently large and (*k*_1_, *k*_2_)_ϵ_^(*i*)^ is the *i*th accepted sample from *p* (*k*_1_, *k*_2_ ∣ *ρ*(**𝒟**(**X**_*s*_), **𝒟**(**X**_*d*_)) < ϵ) using ABC rejection.

Now, consider a geometric sequence of acceptance thresholds ϵ*_ℓ_* = ϵ_0_*K*^-*ℓ*^ for integer *K* ≥ 2 and *ℓ* = 0,1, 2,…,*L* and denote 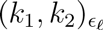 as the random vector distributed according the approximation *p* (*k*_1_, *k*_2_ ∣ *ρ*(**𝒟**(**X**_*s*_), **𝒟**(**X**_*d*_)) < ϵ*_ℓ_*. Using this sequence of posterior distribution approximations, a multilevel estimator for the posterior CDF at the point (*s*_1_, *s*_2_) can be determined as 
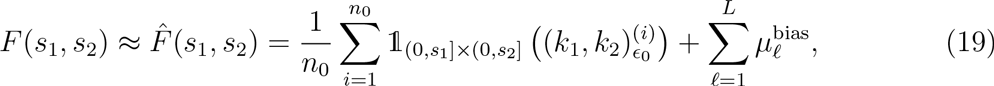
with 
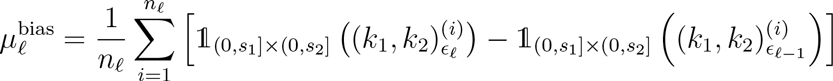
where the sample sizes, *n*_*ℓ*_, are sufficiently large to ensure the estimator variance is below some predetermined value. In terms of bias, the multilevel estimator in Equation (19) is equivalent to the standard Monte Carlo estimator with ϵ = ϵ*_L_* in Equation (18). By evaluating Equation (19) at a set of points in ℝ^2^ combined with a suitable interpolation scheme (see Appendix S3), an approximation of the entire posterior CDF can be constructed.

The estimator given in Equation (19) is the essence of the MLABC method. Although we present the method in terms of the degradation/production model with two unknown parameters, an equivalent general estimator can be formed for any stochastic model with *M* unknown parameters by evaluating the indicator functional over the region (− ∞, *s*_1_] × (− ∞, *s*_2_] ×…× (— ∞, *s*_*m*_]. We now aim to analyze and demonstrate that, under certain reasonable assumptions, this approach can always provide a computational gain over standard ABC rejection sampling for a sufficiently small target bias level.

### Assumptions

There are three main assumptions required for the analysis of Equation (19) [37]. The first two assumptions are related to the convergence rates of the posterior approximation as *ℓ* → ∞TO. The third is a condition on the computation time for generating a sample pair 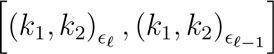 which is required when computing the bias correction term *μ*_ℓ_^bias^. Such a sample pair is a sample from the joint distribution of 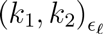 and 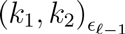.

More specifically, given a geometric sequence of acceptance thresholds ϵ*_ℓ_* = ϵ_0_*K*^-*ℓ*^ for integer *K* ≥ 2, we assume there exist constants *α* > 0, *β* > 0 and γ > 0 such that:

1. as *ℓ* → ∞,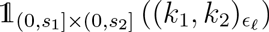 converges weakly to 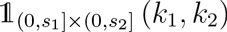 with order α. That is, 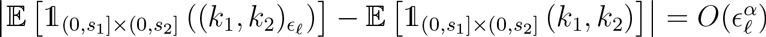
2. as *ℓ* → ∞,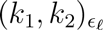 converges strongly to (*k*_1_, *k*_2_) with order *β*. That is, 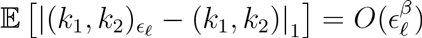.Here, ∣ ․ ∣_1_ is the *ℓ*_1_-norm;
3. the computation time for sampling the distribution of 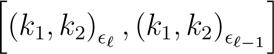 is 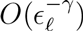

Assumptions 1 and 2 are reasonable due to the convergence properties of ABC rejection itself [18]. However, obtaining exact values or estimates for *α* and *β* is generally difficult. In practice, exact values of *α* and *β* is not required, though we assume these are known in order to analyze the asymptotic computational gain.

The constant *γ* depends on the dimensionality of the parameter space, since the computation time is inversely proportional to the average acceptance rate of the ABC rejection method. The general proof of Assumption 3 is given in Appendix S1. For the degradation/production model we have *γ* = 2.

### Theoretical performance gain

In this section, we present asymptotic bounds on both the RMSE of the standard ABC inference and our MLABC method. We then express the asymptotic computation time given a target RMSE, *h*. This allows us to construct an expression for the asymptotic computational gain in terms of the convergence parameters *α* and β. We do this in the context of a two parameter inference problem, such as the degradation/production model (Equation (10)), leaving more general analysis to Appendix S2.

Assume that the posterior PDF, *p* (*k*_1_, *k*_2_ ∣ **X**_*d*_), has compact support in the region *R* = [*s*_1_^*min*^, *s*_1_^*max*^]×[*s*_2_^*min*^, *s*_2_^*max*^]. Such an assumption is reasonable for a sufficiently large number of observation times *N*_*t*_; for the degradation/production model, this assumption is valid for *N*_*t*_ ≥ 2. Now apply a regular discretization to *R* using *D* divisions in each dimension, resulting in (*D* + 1)^2^ grid points 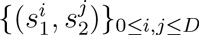, where 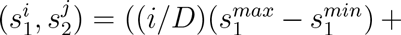 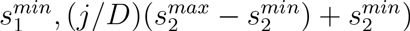.

By applying estimators Equation (18) or Equation (19) we obtain a discrete approximation, 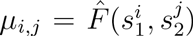, to the true posterior CDF. The RMSE of this approximation is 
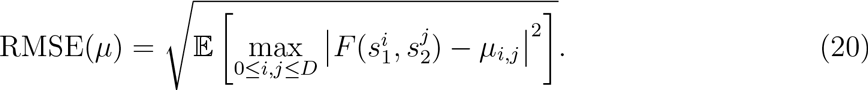

For simplicity, we will ignore the interpolation of *μ*_*i,j*_ for a continuous approximation to *F* over the entire region *R*. It is important to note, however, our analysis (Appendices S2 and S3) accounts for this, and is crucial to our results.

The standard Monte Carlo estimator is given by 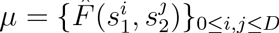 where 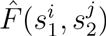 is computed using Equation (18) with ϵ = ϵ*_L_*. If the convergence parameter *α* is known, the RMSE for the standard Monte Carlo estimator, μ, is bounded, for some constant *c*_1_, by 
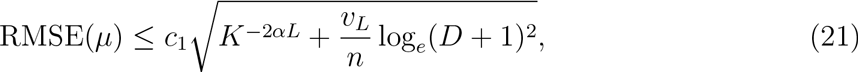
where *υ*_*L*_ is the variance of (*k*_1_, *k*_2_)_ϵ_ and *n* is the number of Monte Carlo samples.

Under Assumption 3, and by taking *L* =(1/*α*) log*_K_*(1/*h*), the cost of using standard Monte Carlo to construct an approximation, *μ*, to the posterior CDF with RMSE(*μ*) = *O*(*h*) is bounded, for some constant *c*_2_, by 
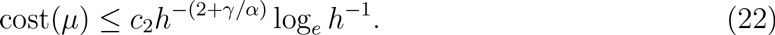

This bound is identified by bounding the right hand side of Equation (20) by *h* and solving for a lower bound on the number of accepted samples, *n*.

Bounding the RMSE for the multilevel estimator, 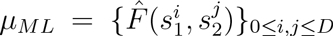 with 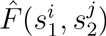 computed using Equation (19), is a relatively straightforward application of the method for obtaining Equation (20) along with invoking Assumption 2. The equivalent result for the multilevel estimator, for some constant *c*_3_, is 
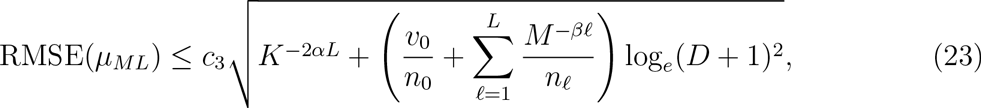
where *υ*_0_ is the variance of (*k*_1_, *k*_2_)_ϵ0_ and *n_ℓ_* are the Monte Carlo sample numbers for each level.

The upper bound on the computation time for *μ*_*ML*_ depends on the choice of the number of samples for each level, *n_l_*. We can use a Lagrange multiplier method to choose the *n*_*ℓ*_ such that the asymptotic cost is minimized under the constraint of RMSE(*μ*_*ML*_) = *O*(*h*). The choice of *n_ℓ_* is 
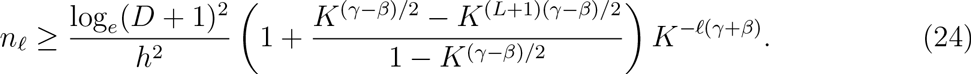

Using Assumption 3 and the optimal *n*_*ℓ*_ given by Equation (24), we arrive at an expression for the asymptotic bound on the computation time for evaluating *μ*_*ML*_. In the case of *β* < *γ*, then there exists some constant *c*_4_ such that 
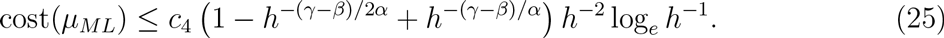

The results from Equation (22) and Equation (25) directly yield a reduction in the order of the computation time upper bound from using the multilevel method. We denote the reduction as the asymptotic computational gain, given by 
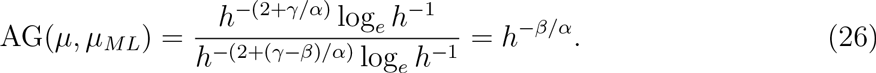

Note that Equation (26) indicates that, assuming the upper bounds are achieved asymptotically, it is always possible to choose a target RMSE, *h*, such that the multilevel approach achieves some desired level of computational gain. The rate of growth of this gain as a function of h depends on the convergence characteristics of the sequence of ABC posterior approximations, however it is always an improvement since *α* > 0 and *β* > 0.

## Practical application of MLABC

In practice, the convergence parameters *α* and *β* are unknown. In this section, we present a practical implementation of an algorithm for MLABC that does not depend on explicit knowledge of *α* and β. We find that our algorithm can obtain up to an order of magnitude performance improvement while still maintaining a desired level of accuracy.

### Removing dependence on convergence parameters

First, note that when ABC methods such as ABC rejection are used in practice, there are certain assumptions made about the weak convergence rate, α. If we expect *n*— accepted samples using acceptance threshold ϵ to provide an acceptable estimator of the real posterior CDF, then we are implicitly assuming 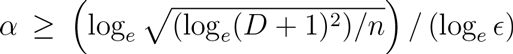. Therefore, we can determine, for a given scale factor, *K*, and base level threshold, ϵ_0_, the number of levels, *L*, required to match the final bias of the standard ABC rejection method. As a result, we can treat *α* as a known parameter in the sense of equivalence to the standard ABC rejection estimator.

Second, consider the role of *β* in determining the *n_ℓ_* in Equation (24). While the details are given in Appendix S2, we informally state that the summation in Equation (24) can be derived by showing that Assumption 2 implies, for some constant *c*_5_, 
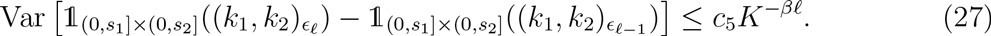

That is, *K*^-*βℓ*^ is used to bound the variances of the bias correction terms in Equation (19). It is important to note that the rigorous version of Equation (27) requires a smoothing process to be applied to the indicator functional to ensure the Lipschitz continuity conditions discussed in the supporting information (Appendix S2 and S3).

If we knew exactly, for each level *ℓ*, the computation time to generate a posterior realization, *c*_*ℓ*_, and the variance, *υ*_*ℓ*_, of each bias correction term, *μ*_*ℓ*_^*bias*^, then we could calculate the optimal *n*_*ℓ*_ using the same Lagrange multiplier approach as for Equation (24).

**Algorithm 3.**
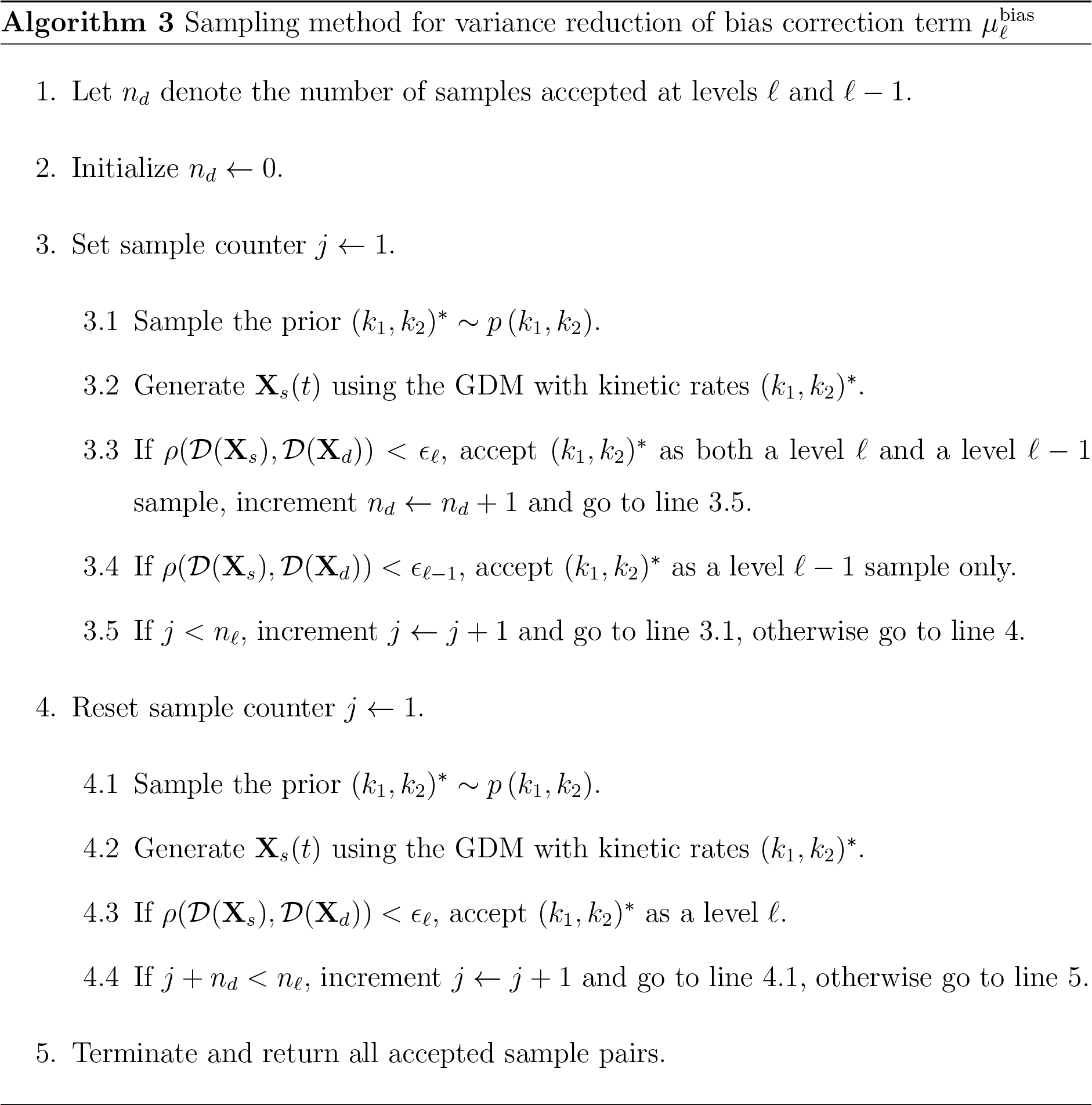

The result would be, 
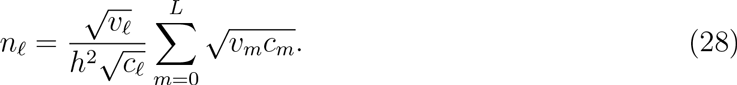

In practice we can can only estimate *c*_*ℓ*_ and *υ*_*ℓ*_. Using the approach used by Anderson et al. [33] and Lester et al. [34] we generate a relatively small number of trial samples at each level to obtain estimates for *c*_*ℓ*_ and *υ*_*ℓ*_ that work well in practice. Using this approach we do not have the same theoretical bounds on the RMSE. However, accurate approximations for the variances will result in estimators with a RMSE that is close to the target in practice.

**Algorithm 4.**
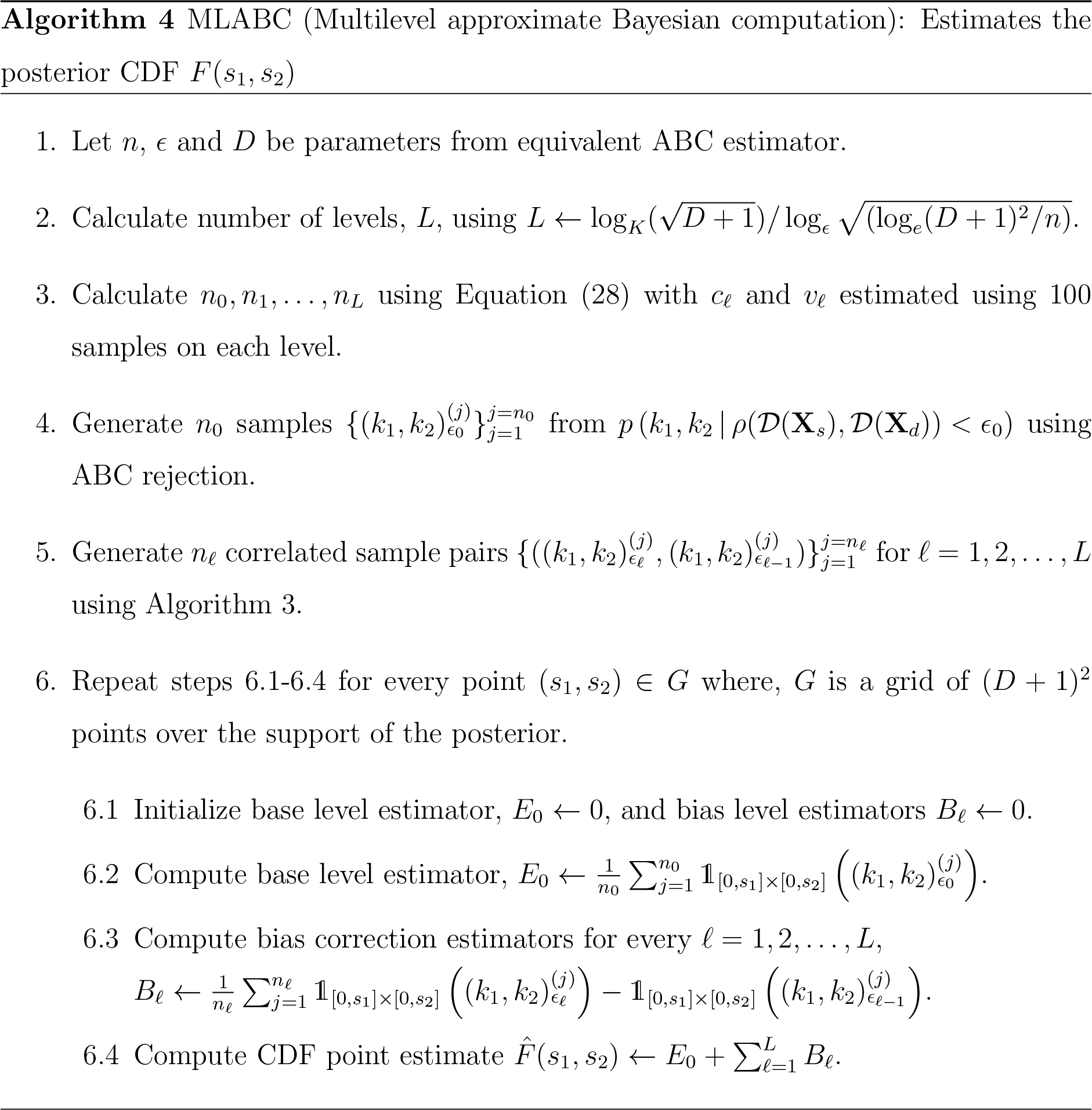

### Improving performance with variance reduction

In Equation (28), note that 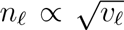Furthermore, note that for the bias correction terms in Equation (19), our only concern is the expected difference between indicator function values at each level rather than the expected values themselves. As a result, if we can introduce some correlation in the generation of accepted samples at each level, then the variance of the estimators and hence the number of samples required will decrease. In this work, we implement a simple method to ensure such correlation. Given a simulated trajectory, **X**_*s*_(*t*), then *ρ*(**𝒟**(**X**_*s*_),**𝒟**(**X**_*d*_)) < ϵ*_ℓ_* implies *ρ*(**𝒟**(**X**_*s*_),**𝒟**(**X**_*d*_)) < ϵ_*ℓ*-1_, since ϵ*_ℓ_* < ϵ_*ℓ*-1_. As a result, we can sample level *ℓ* — 1 first and keep track of any samples that can also be accepted at level *ℓ*; such samples can be validly taken as samples from both levels. A pair constructed this way will make no contribution to the bias correction term, hence reducing the variance. We arrive at the method presented in Algorithm 3 for reducing the variance when sampling for the bias correction term μ_*ℓ*_^bias^.

### The multilevel ABC algorithm

By combining Equation (28) with Algorithm 3, we construct a practical implementation of the MLABC estimator (Equation (19)). This MLABC method, presented in Algorithm 4 for the degradation/production model, is implemented as a prototype using the C programming language. The prototype (Code S3) is specific to biochemical reaction network models, however, only minor changes are required to change the target application.

## Results

In this section, we examine the accuracy and performance of MLABC using the previously presented degradation and degradation/production models because we can directly evaluate the estimator RMSE using the exact solution to the CME. We then compare MLABC with standard ABC rejection using two more complex biochemical reaction networks.

### Estimating convergence parameters

The asymptotic computational gain using MLABC is presented in Equation (26). To validate this theoretical result, we need to estimate *α* and *β*. We use the ABC rejection method (Algorithm 2) and linear regression to estimate *α* and *β* for the degradation model and the degradation/production model using data observed at *N*_*t*_ discrete points in time for *N*_*t*_ = 2, 3,…, 10. For the degradation model and degradation/production model, respectively, data is generated using *k* = 0.1 and (*k*_1_, *k*_2_) = (0.1,1). A univariate uniform prior distribution with support {*k*: *k* ∊ [0,1]} is utilized for the degradation model and a bivariate uniform prior with support {(*k*_1_, *k*_2_): (*k*_1_, *k*_2_) ∊ [0,1] × [0,10]} is utilized for the degradation/production model. Estimates are produced using 1, 000 accepted samples for threshold levels ϵ*_l_* = ϵ_0_*K*^-*ℓ*^, *ℓ* = 1, 2,…, *L* with *K* = 2, ϵ_0_ = 1 and *L* = 5. These estimates are shown in Figure 3(a)-(b).

**Figure 3.**
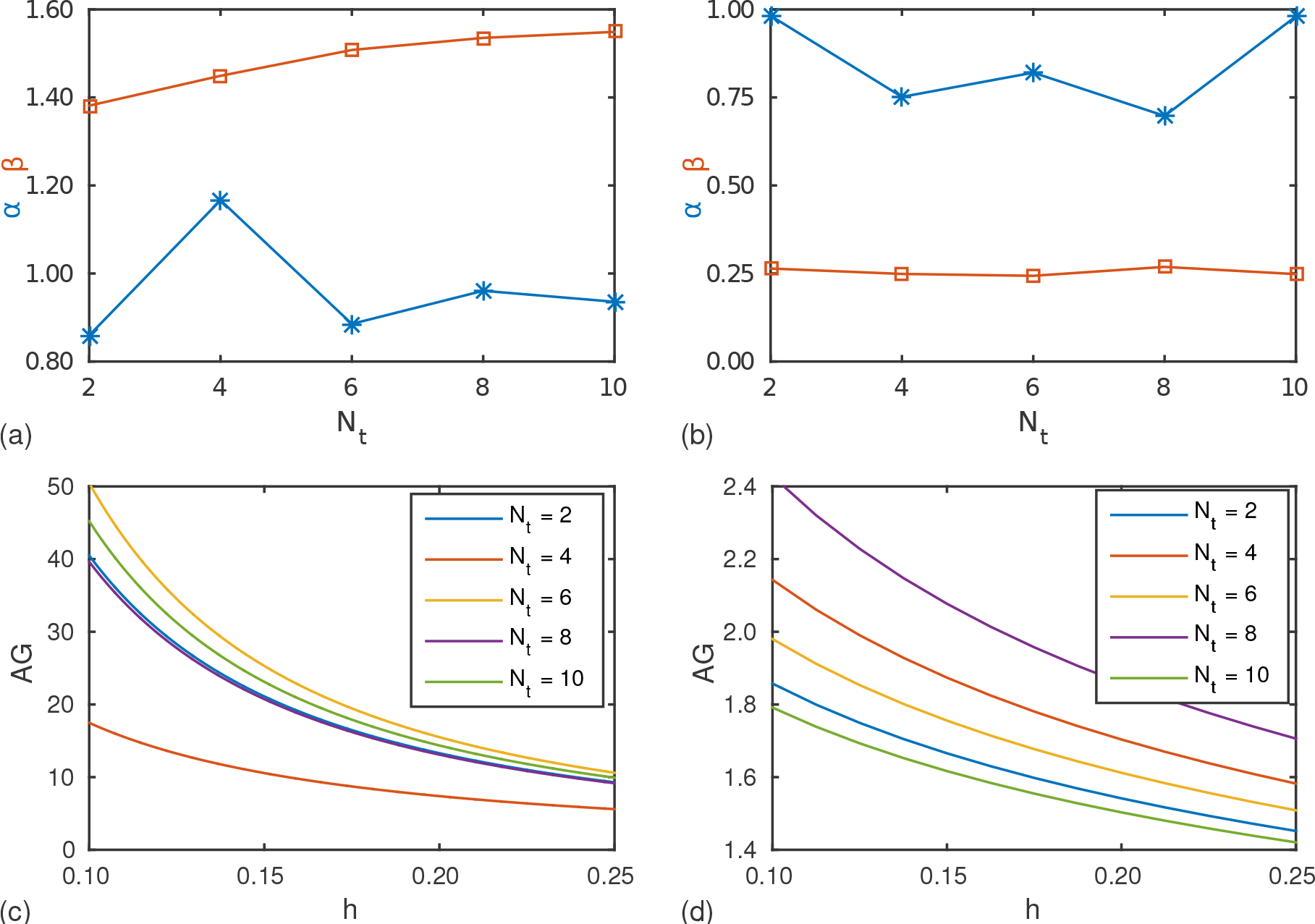
Estimated convergence rates. Least squares approximation of *α* and *β* for the (a) degradation model with *k* = 0.1 and (b) degradation/production model with (*k*_1_, *k*_2_) = (0.1,1) using observation times *N*_*t*_ ∊ [2,…, 10]. Asymptotic computational gain functions *AG* = *O*(*h*^-*β*/*α*^) for the (c) degradation and (d) degradation/production models.

Using these estimates of *α* and *β* it is possible to predict the asymptotic growth of computational gain by using MLABC based on sample numbers chosen according to Equation (24). These computational gains are given in Figure 3(c)-(d) for *N*_*t*_ = 2, 3,…, 10. By using Equation (24), we expect minimal increase in the RMSE compared with standard ABC rejection.

### Performance using estimated convergence parameters

We now compare numerical simulation results against the theoretical performance and error analysis. Throughout we refer to the measured computational gain of our MLABC approach that is given by 
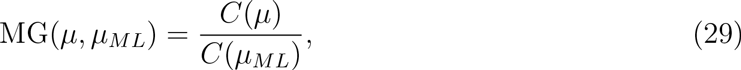
where *μ* and *μ*_*ML*_ are the standard ABC rejection and MLABC estimators, respectively, and *C*(·) is the averaged measured computation time to evaluate the estimator.

For the degradation model, 20 independent MLABC and ABC rejection CDF estimators are computed for target RMSE *h* ∊ [0.1, 0.25]. The CDF is approximated over the interval [0, 0.5] on a grid of 1, 000 nodes using data with *N*_*t*_ = 8. The sample numbers, *n*_*ℓ*_, are computed according to Equation (24) using the estimated convergence parameter values of *α* ≈ 0.96 and *β* ≈ 1.54. As shown in Figure 4(a), the increase in computational gain (Equation (29)) is the same order of magnitude as the theoretical asymptotic prediction (Equation (26)), albeit at a smaller absolute scale. In Figure 4(b), we see that the target of RMSE ≤ *h* is achieved with the exception of *h* = 0.1, however, the fact that there is also an increase in RMSE at *h* = 0.1 for the standard ABC estimator indicates that this is a feature of the problem that can be probably be improved with the application of smoothing to the indicator functional (see Appendix S3).

For the degradation/production model, 20 independent MLABC and ABC rejection CDF estimators are computed for target RMSE *h* ∊ [0.1, 0.25]. The CDF is approximated over the region [0,1] × [0,10] on a grid of 100 × 100 nodes using data with *N*_*t*_ = 4. The sample numbers, *n*_*l*_, are computed according to Equation (24) using the estimated convergence parameter values of *α* ≈ 0.75 and *β* ≈ 0.25. Figure 4(c)-(d) provides, for the production/degradation model, the computational gain using MLABC over ABC rejection and the RMSE versus the target RMSE. Compared to the degradation model, the computational gain is significantly less; however, it is consistent with the theoretical results. Similarly, Figure 4(d) shows practically no increase in the RMSE of the CDF estimator.

An important remark must be made about these results. They do not represent the best computational gain that can be achieved in practice, but rather they provide experimental validation for our theory. If the values of a and are known, then it is possible to predict the asymptotic computational gain available whilst maintaining control on the RMSE of the estimator. Furthermore, we note that, as predicted by the theory, the computational gain grows proportionally to a power of *h*^-1^.

**Figure 4.**
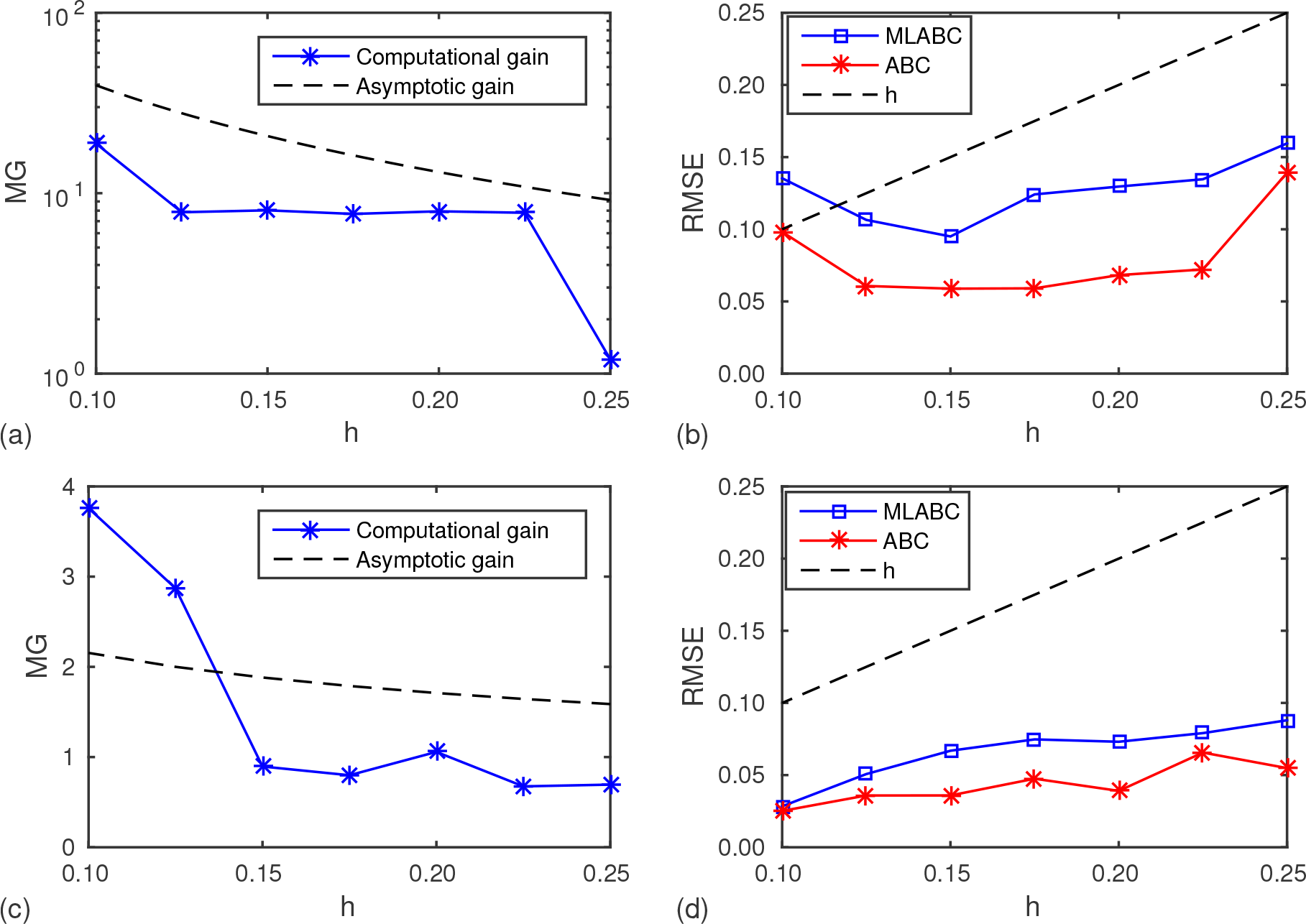
Measured performance gain and error. (a) Computational gain and
(b) RMSE for the degradation model results using data with *N*_*t*_ = 8 observation times and convergence rate estimates *α* ≈ 0.96 and *β* ≈ 1.54. (c) Computational gain and (d) RMSE for the degradation/production model results using data with *N*_*t*_ =4 observation times and convergence rate estimates *α* ≈ 0.75 and *β* ≈ 0.25.

### Performance using empirical sample numbers

We now look to the more practical approach to MLABC, as presented in Algorithm 4. Here we specifically focus on the degradation/production model. In the first experiment, 20 independent MLABC (using Algorithm 4) and ABC rejection CDF estimators of the degradation/production model are computed for target RMSE *h* ∊ [0.1, 0.25]. The CDF is approximated over the support region {(*k*_1_, *k*_2_): (*k*_1_, *k*_2_) ∊ [0,1] × [0,10]} on a grid of 100 × 100 nodes using data with *N*_*t*_ = 4.

Results in Figure 5 are analogous to the results in Figure 4(c)-(d). By using Algorithm 4, we have achieved greater computational gain, whilst maintaining reasonable control over the RMSE (i.e., still within target *h*). The main advantage is that values for a and have not been required. While the performance results in Figure 5(a) are an improvement over those in Figure 4(c), the new results still show the same order of magnitude increase in performance. However, we now demonstrate that Algorithm 4 can out perform the asymptotic results by an order of magnitude.

**Figure 5.**
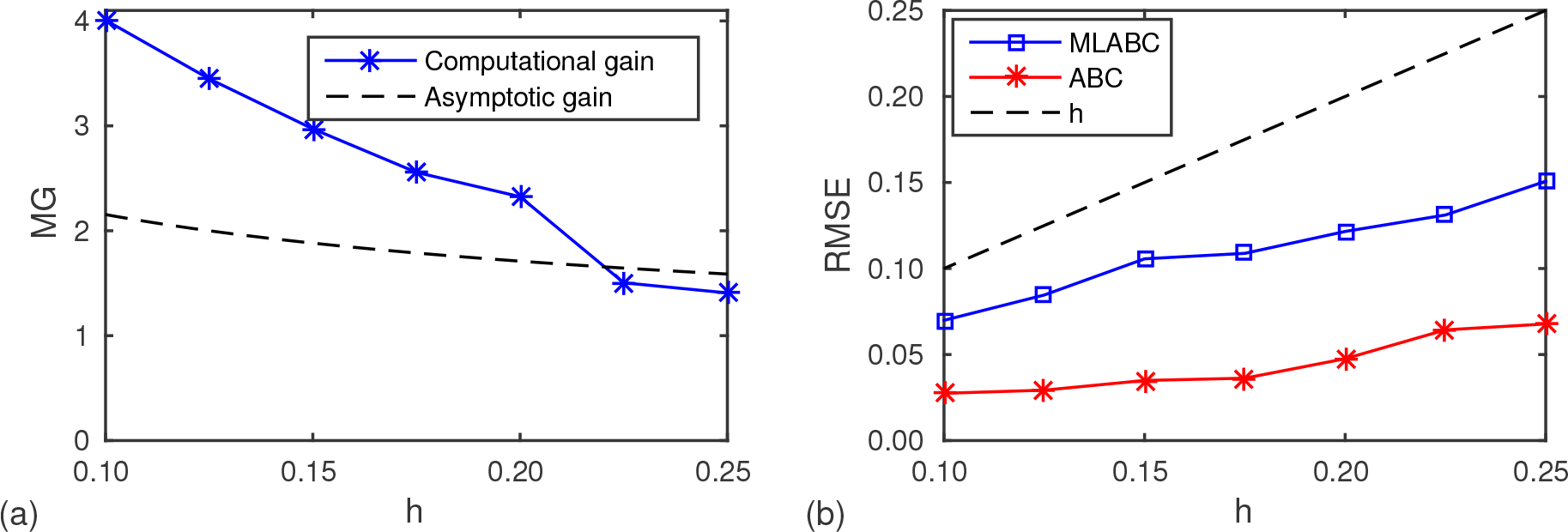
Measured performance gain and error for the degradation/production model. Using data with *N*_*t*_ = 4 observation times and 100 samples at each level to select sample numbers *n*_*ℓ*_. (a) Measured computational gain and (b) RMSE as a functions of *h*.

In the second experiment, the target RMSE is kept constant at *h* = 0.2. MLABC and ABC rejection estimators for the posterior CDF are computed using data with *N*_*t*_ = 2,4,6,…, 20. For each value of *N*_*t*_, 20 simulations are executed, with computation times being the average. Figure 6(a) demonstrates a significant improvement in computation time for MLABC over ABC rejection in this case. Note that the computational gain shown in Figure 6(b) is an order of magnitude greater than the asymptotic analysis shown in Figure 3(d) predicts for *N*_*t*_ = 10.

We now compare the quality of the posteriors computed by MLABC and ABC rejection. For this we consider the posterior mean and the marginal distribution 90% credible intervals. The mean, representing the central tendency of the posterior, represents a likely parameter candidate, (*k*_1_^*m*^, *k*_2_^*m*^), given by 
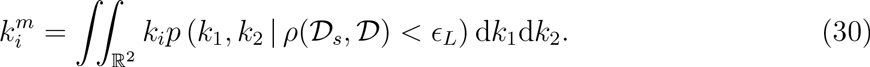

**Figure 6.**
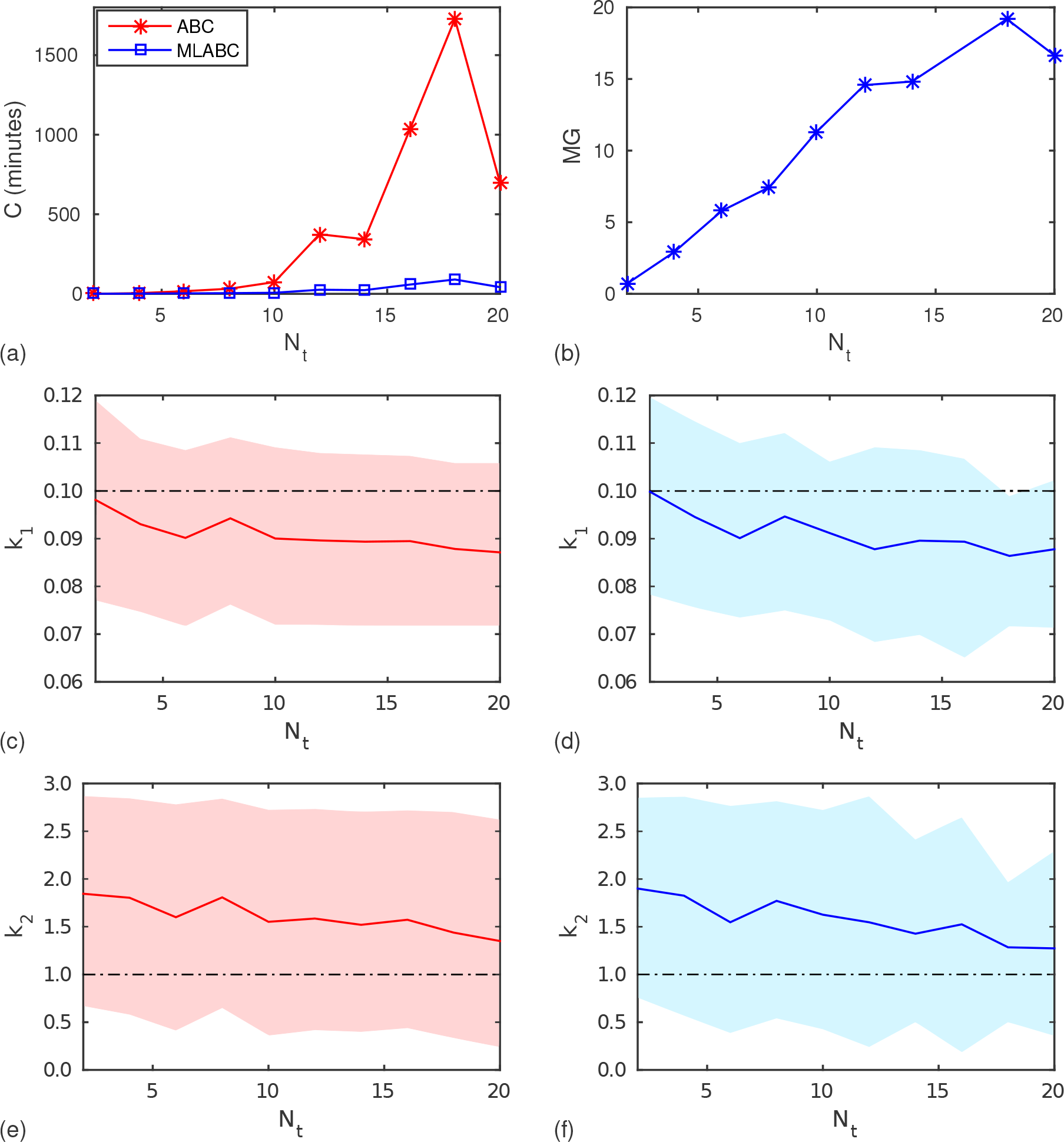
Comparison of ABC and MLABC: degradation/production. Performance of ABC and MLABC for the degradation/production model as *N*_*t*_ increases. (a) Computation time. (b) Computational gain. (c) and (e) ABC parameter estimates. (d) and (f) MLABC parameter estimates. The true parameter values, (*k*_1_, *k*_2_) = (0.1,1.0), are indicated with dashed lines, posterior means and 90% credible intervals are indicated with solid lines and shaded areas, respectively.

Given the joint CDF, *F*(*s*_1_, *s*_2_), the marginal CDFs, *F*_1_(*s*) and *F*_2_(*s*), can be determined using 
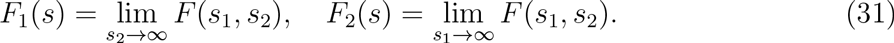

For some significance level, *a*, the (1 — *a*)100% credible interval for parameter *k*_*i*_ denoted by [*ℓ*(*k*_*i*_),*u*(*k*_*i*_)], is 
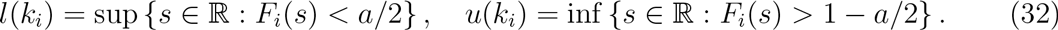

The credible interval provides a measure of uncertainty around the parameter estimates.

Figures 6(c) and 6(e) present the estimates produced by standard ABC for different values of *N*_*t*_. These are to be compared with the estimates produced by MLABC as shown in Figures 6(d) and 6(f). These results show that, from a practical perspective, the MLABC method is just as appropriate as ABC rejection. That is, the true parameter values lie within the credible region of the posteriors and these credible intervals are almost the same for MLABC compared with ABC rejection. Both methods yield very similar mean and credible interval values.

### Higher dimensional and higher order models

We now investigate the validity of our MLABC approach to inference for higher dimensional and higher order models. In the first instance, we investigate the inference problem in four-dimensional parameter spaces for first order reactions. We then investigate a more biologically inspired enzyme kinetics model that includes a second order reaction.

#### Mono-molecular chains

A general mono-molecular chain biochemical reaction network has the form 
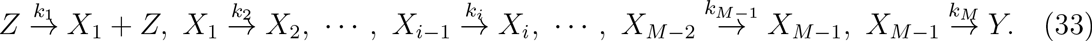

As shown by Jahnke et al. [41], the CME for Equation (33) has a closed form solution, however, it is non-trivial to evaluate. For the purposes of this study, we suppose the CME of the mono-molecular chain to be intractable.

When *M* = 4 we have the four reaction mono-molecular chain 
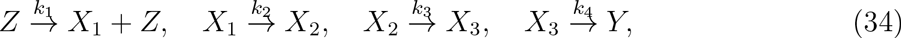
with initial conditions 
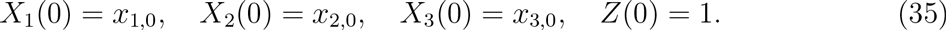

We compute 20 independent MLABC and ABC rejection CDF estimators for the four reaction mono-molecular chains (Equation (34)) using data with *N*_*t*_ = [2, 4,…, 20] observation points. The CDFs are approximated over the region {(*k*_1_ *k*_2_, *k*_3_, *k*_4_): (*k*_1_, *k*_2_, *k*_3_, *k*_4_) ∊ [0, 3] × [0, 0.3] × [0, 0.03] × [0, 0.03]} using the target RMSE *h* = 0.2. Computation times are averaged over the 20 samples.

For the four reaction mono-molecular chain model, we note in Figure 7(b) that a peak computational gain of approximately five times is achieved before a reduction back:o four times. Figures 7(c)-(j) also provides the resulting parameter estimates using ABC and MLABC. In this case, the MLABC and standard ABC estimates are in very close agreement across all four parameters. The MLABC estimator displays more variability in *u*(*k*_*i*_) for *N*_*t*_ ≥ 10, however these oscillations follow the same trend as the ABC estimates. We note that for a three reaction mono-molecular chain the results (not shown) are very similar, however, a peak of 10 times computational gain is observed.

One final remark on the results for mono-molecular chains: note that the degrada-ion/production model is actually a mono-molecular chain with *M* = 2. In light of this, it is interesting to note that the peak computational gain achieved for the degrada-:ion/production model in Figure 6(b) is around 20 times (*M* = 2), for the three reaction mono-molecular chain it is around 10 times (*M* = 3)(results not shown) and for the bur reaction mono-molecular chain it is around five times (*M* = 4) (Figure 7(b)). This could indicate that the ratio of the convergence rates, *β*/*α*, is inversely proportional to the number of reactions, *M*. There could be other factors at work here, but given the curse of dimensionality inversely affects the order of the acceptance rate, *γ*, it is logical to conclude that there is also an additional influence on the convergence rates *α* and *β*. More experimental and theoretical work is needed to analyze the relationship between these convergence rates and the dimensionality of the parameter space.

**Figure 7.**
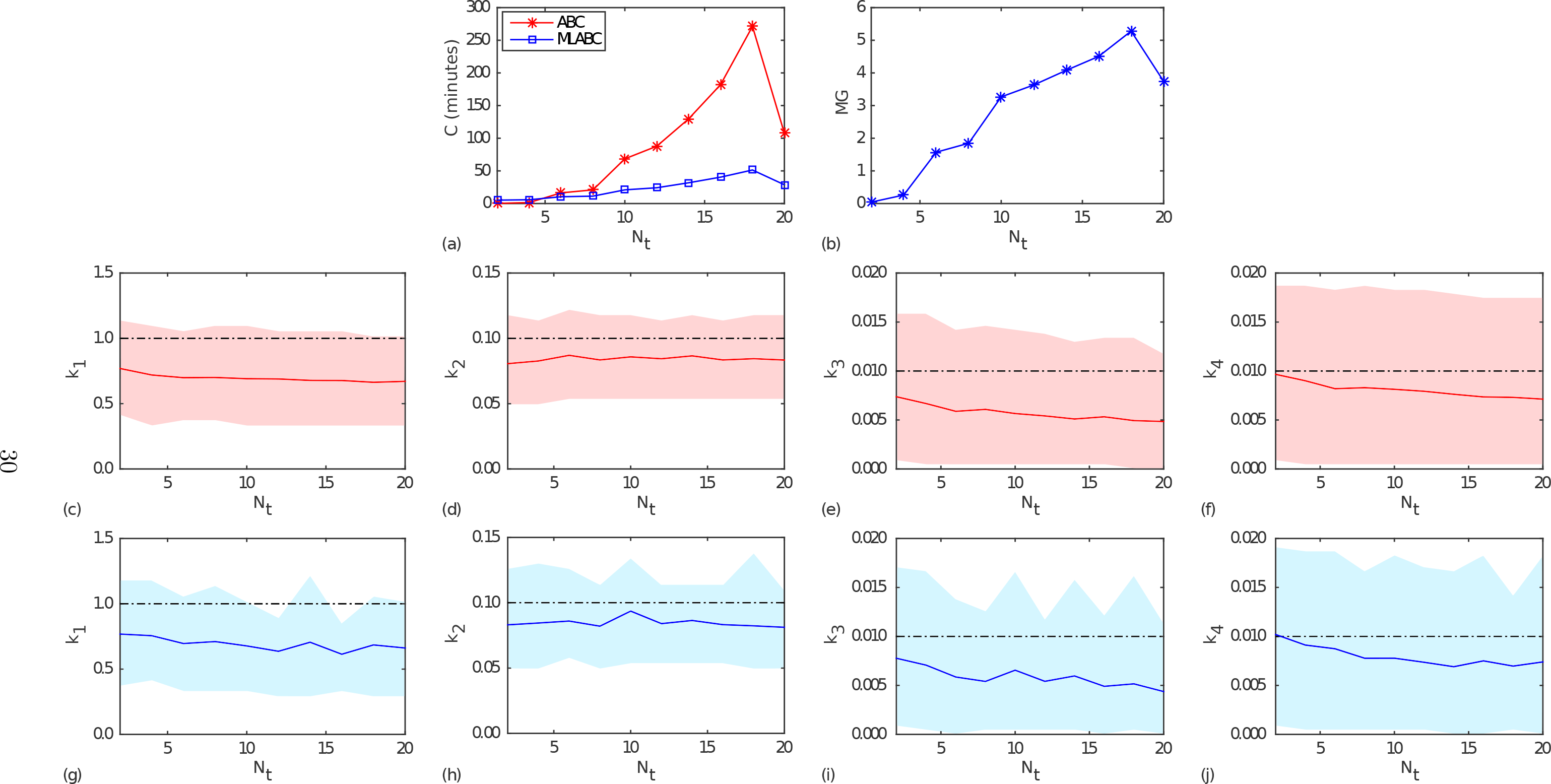
Comparison of ABC and MLABC: Four reaction mono-molecular chain. Performance of ABC and MLABC for the four reaction mono-molecular chain as the number of observation times, *N*_*t*_, increases. (a) Computation time. (b) Computational gain. (c)-(f) ABC parameter estimates. (g)-(j) MLABC parameter estimates. The true parameter values, (*k*_1_, *k*_2_, *k*_3_, *k*_4_) = (1, 0.1, 0.01, 0.01), are indicated with dashed lines, posterior means and 90% credible intervals are indicated with solid lines and shaded areas, respectively.

#### Higher order models

We now test the applicability of MLABC to more general biochemical reaction networks with higher order reactions. Such networks rarely yield a tractable solution to the CME [7, 39]. As a result, such higher order models are practical target applications for MLABC. We consider a Michaelis-Menten enzyme kinetic model [42], which describes the dynamics of an enzyme-catalyzed reaction of a substrate *S* into a product *P* with the enzyme *E* acting as a catalyst. A three reaction Michaelis-Menten model is given by 
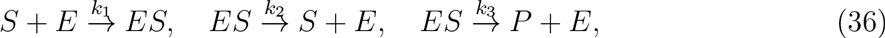
with initial conditions 
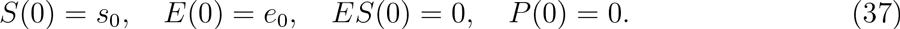

An example realization demonstrating the additional complexity of the dynamics of this model is provided in Figure 8. The realization displays the conversion of *S* into *P*. Note that as *S*(*t*) → (*k*_2_ + *k*_3_/*k*_1_), the propensity function of the third reaction *a*_3_(*ES*(*t*); *k*_3_) → *k*_3_*e*_0_/2, so the shape of the *ES* curve depends crucially on this ratio of parameters. As a result, more observations time points are required to ensure the assumption of compact support for the posterior is reasonable.

**Figure 8.**
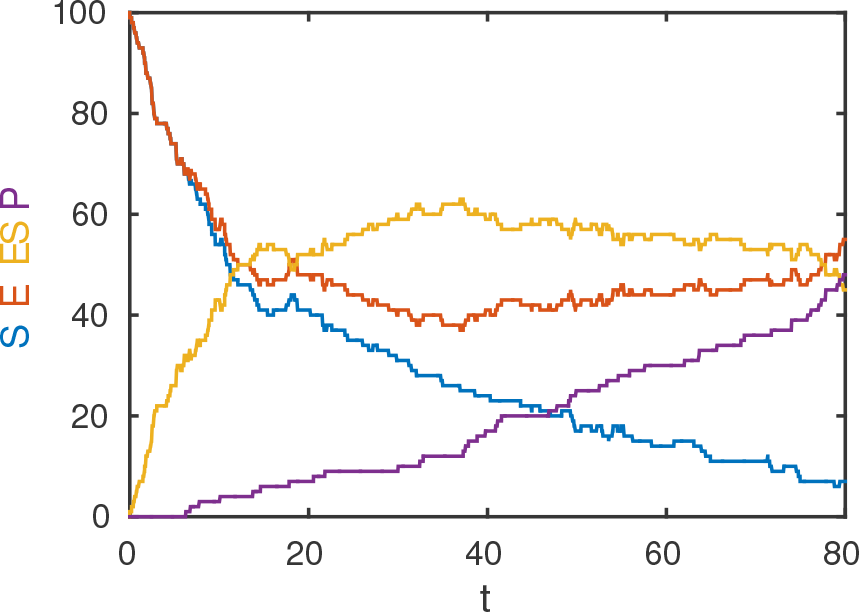
Michaelis-Menten model realization. Example realization with *k*_1_ = 0.001 (sec^-1^), *k*_2_ = 0.005 (sec^-1^), *k*_3_ = 0.01 (sec^-1^) and *s*_0_ = *e*_0_ = 100.

The Michaelis-Menten model given in Equation (36) is the first we investigate with an intractable CME. As a result we investigate how the performance of MLABC is affected by both the choice of target RMSE, *h*, and the choice of the number of observation points, *N*_*t*_.

First, we consider the computational gain achieved for the inference on the Michaelis-Menten model as the target RMSE, *h*, decreases. This time, the CDF estimators are constructed using ABC and MLABC with a uniform prior with support {(*k*_1_, *k*_2_, *k*_3_: (*k*_1_, *k*_2_, *k*_3_) ∊ [0, 0.003] × [0, 0.015] × [0, 0.03]} and fixed number of observation point, *N*_*t*_ =
12. Estimators are computed for target RMSEs of *h* = 0.125, 0.15,…, 0.2. Computation times are averaged over 20 independent simulations. Results are shown in Figure 9, confirming the growth in computational gain as *h* → 0.

**Figure 9.**
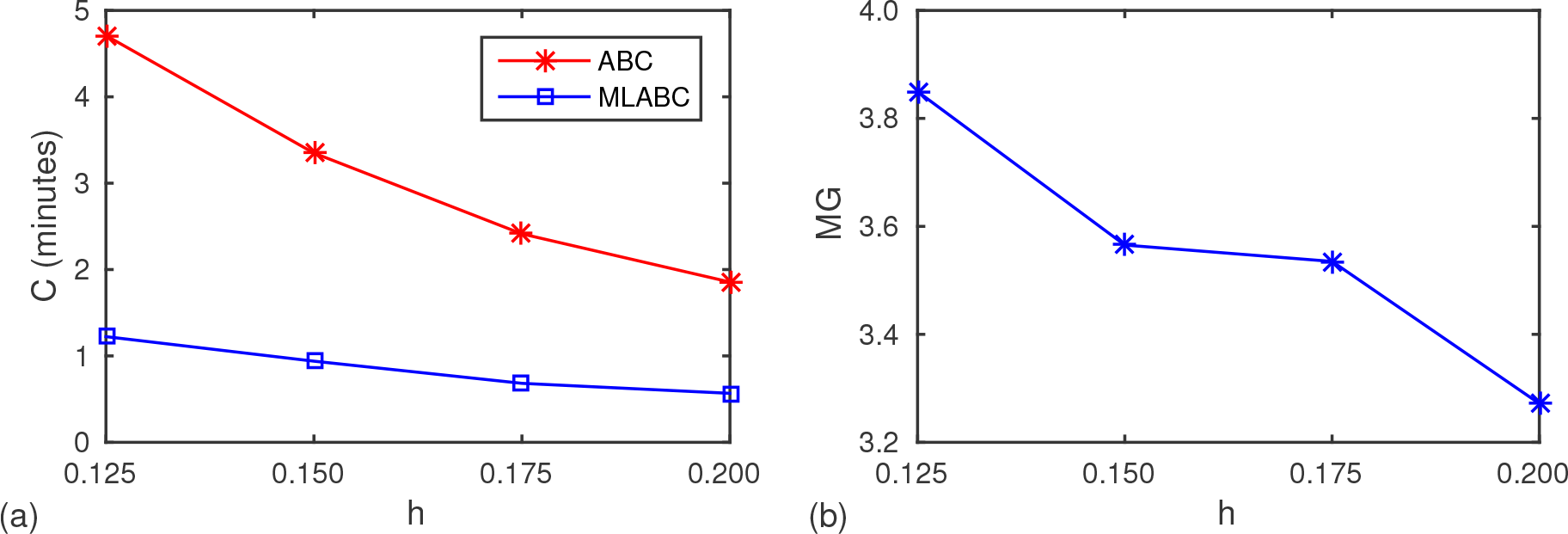
Comparison of ABC and MLABC: Michaelis-Menten model. The performance of MLABC as the target error h decreases. (a) Computation time. (b) Computational gain.

For our last experiment, we take data over a much larger interval *N*_*t*_ = [2,12,… 192]. We choose this interval to highlight the fact that as *N*_*t*_ is increased the computational gain reaches a maximum then plateaus. We conjecture that this reflects a point at which little information is gained through the additional observations. Just as with our other models we compute 20 independent CDF estimators using ABC and MLABC for each value of *N*_*t*_ with target RMSE *h* = 0.2. In this case, we take a uniform prior with support {(*k*_1_, *k*_2_, *k*_3_): (*k*_1_, *k*_2_, *k*_3_) ∊ [0, 0.005] × [0, 0.025] × [0, 0.05]}. The result is a peak computational gain of six times followed by a plateau between four and six times, as shown in Figure 10(b).

The associated parameter estimates are given in Figure 10(c)-(h). The MLABC estimator follows very closely the ABC estimator in terms of the mean for all parameters however the uncertainties in the MLABC estimate for *k*_2_ are nearly double that of the ABC estimator for some values of *N*_*t*_. Interestingly, this seems to occur for values of *N*_*t*_

**Figure 10.**
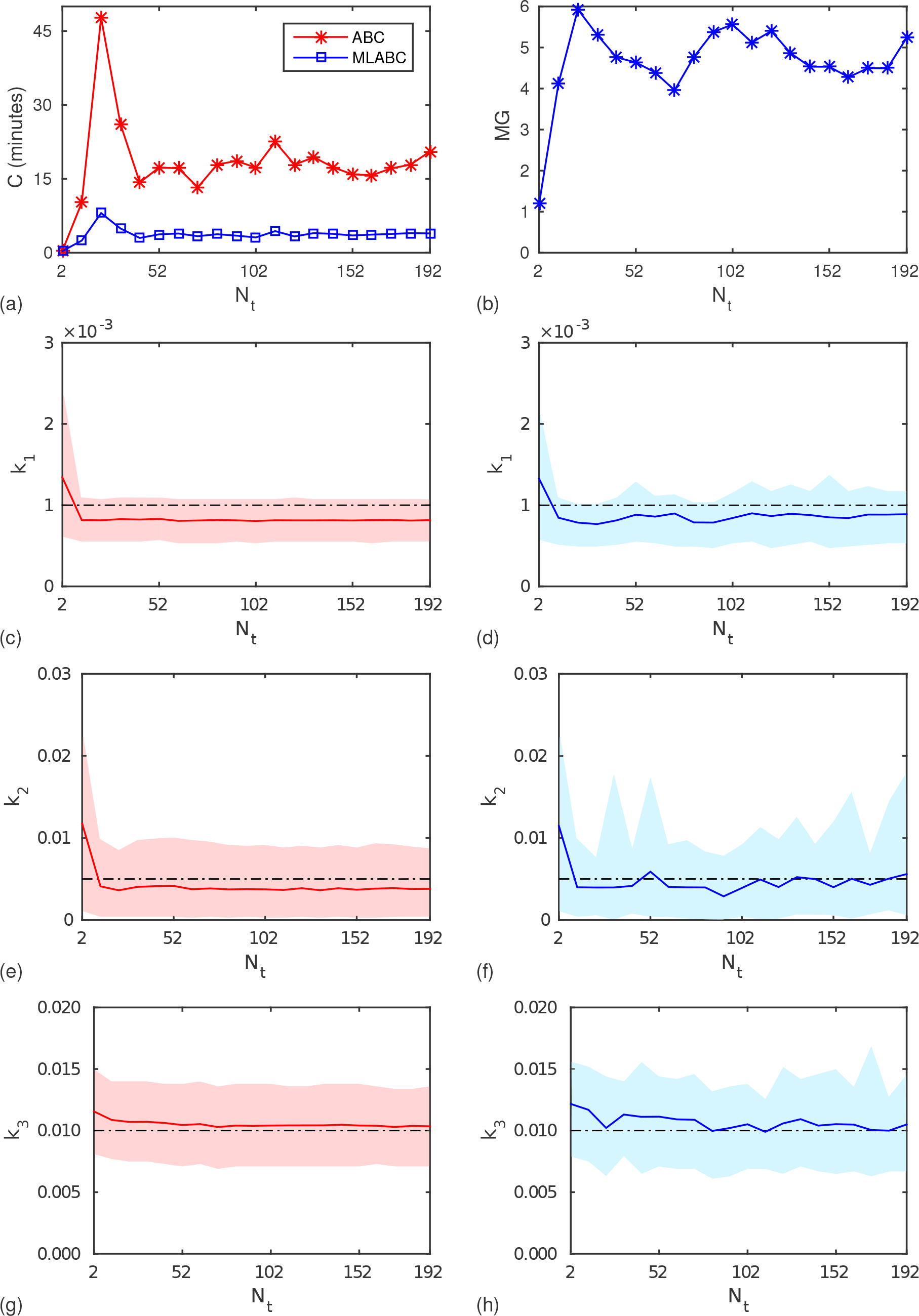
Comparison of ABC and MLABC: Michaelis-Menten model. Performance of ABC and MLABC for the Michaelis-Menten model as the number of observation times, *N*_*t*_, increases. (a) Computation time. (b) Computational gain. (c), (e) and (g) ABC parameter estimates. (d), (f) and (h) MLABC parameter estimates. The true parameter values, (*k*_1_, *k*_2_, *k*_3_) = (0.005, 0.025, 0.05), are indicated with dashed lines, posterior means and 90% credible intervals are indicated with solid lines and in which the computational gain is lower. Further investigation is required to explain the reasons for this.

## Conclusion

In this study, we present a new approach to computational Bayesian inference using MLMC sampling. We perform a general analysis based on the approximation of posterior CDFs using MLMC techniques developed by Giles et al. [37], to show that under our convergence assumptions, asymptotically, a net computational gain is always achievable for some sufficiently small value of RMSE, *h*, and simulation results confirm this prediction.

We also develop a practical implementation of the MLABC method that does not require the convergence rate parameters to be known *a priori*. Numerical estimates of the posterior CDF over a range of models suggest that a computational gain of four to 20 times can often be achieved over standard ABC rejection, with larger data set dimensionality improving this gain, up to some maximum. Under the right conditions, such as one-dimensional inference problems, a computational gain of up to 60 times is achievable.

Though the target application of this work is parameter inference for stochastic biochemical reaction network models, the MLABC method is as general as ABC rejection. From a practical perspective, MLABC can be used in place of ABC rejection if the standard assumptions on weak and strong convergence hold. Minor modifications to the provided prototype code are required to achieve this.

While our current approach is a step towards dealing with the curse of dimensionality, there is still much work to be done. For the purposes of this initial investigation, we use ABC rejection for our benchmark inference method and as the basis of the MLABC method. The most natural extension of this work is the application of the MLMC framework to other ABC methods. We acknowledge that more advanced approaches such *likelihood-free Markov chain Monte Carlo* [26] and *likelihood-free sequential Monte Carlo* [27] will generally deal with higher dimensional models and data more efficiently than ABC rejection. Our MLABC framework, however, is sufficiently general that it is not intimately tied to ABC rejection and there will be future opportunities to apply this approach to these more advanced methods.

## Acknowledgments

This project utilized the high performance computing (HPC) facility at the Queensland University of Technology (QUT). We thank Mike Giles for useful advice.

## Supporting Information

**S1. Derivation of average acceptance rate.**

**S2. Multilevel ABC asymptotic performance analysis.**

**S3. Smoothing and extension.**

**S3. MLABC prototype code.**

